# Identification of community-consensus clinically relevant variants and development of single molecule molecular inversion probes using the CIViC database

**DOI:** 10.1101/479394

**Authors:** Erica K. Barnell, Adam Waalkes, Kelsi Penewit, Katie M. Campbell, Zachary L. Skidmore, Colin C. Pritchard, Todd A. Fehniger, Ravindra Uppaluri, Ramaswamy Govindan, Malachi Griffith, Stephen J. Salipante, Obi L. Griffith

## Abstract

Clinical targeted sequencing panels are important for identifying actionable variants for cancer patients, however, there are currently no strategies to create impartial and rationally-designed panels to accommodate rapidly growing knowledge within the field. Here we use the Clinical Interpretations of Variants in Cancer database (CIViC) in conjunction with single-molecule molecular inversion probe (smMIP) capture to identify and design probes targeting clinically relevant variants in cancer. In total, 2,027 smMIPs were designed to target 111 eligible CIViC variants. The total genomic region covered by the CIViC smMIPs reagent was 61.5 kb. When compared to existing genome or exome sequencing results (n = 27 cancer samples from 5 tumor types), CIViC smMIP sequencing demonstrated a 95% sensitivity for variant detection (n = 61/64 variants). Variant allele frequency for variants identified on both sequencing platforms were highly concordant (Pearson correlation = 0.885; n = 61 variants). Moreover, for individuals with paired tumor/normal samples (n = 12), 182 clinically relevant variants missed by original sequencing were discovered by CIViC smMIPs sequencing. This design paradigm demonstrates the utility of an open-sourced database built on attendant community contributions for each variant with peer-reviewed interpretations. Use of a public repository for variant identification, probe development, and variant annotation could provide a transparent approach to build a dynamic next-generation sequencing–based oncology panel.

## Introduction

Despite recognition that genomics plays an important role in tumor prognosis, diagnosis, and treatment, scaling genetic analysis to encompass every clinical tumor specimen has been unattainable.^1,2^ Barriers preventing widespread adoption of genomic analysis into treatment protocols include: costs associated with genomic sequencing and analysis,^3^ computational limitations preventing timely and accurate annotation of variants,^3^ and extensive growth of knowledge within the precision oncology field.^4^ Technological improvements in sequencing and data analysis continue to reduce these first two limitations, however, less progress has been made in integrating dynamic genomic annotation into clinical diagnoses. Over 22% of oncologists have acknowledged limited confidence in their own understanding of how genomic knowledge applies to patients’ treatment and 18% reported testing patients’ genetics infrequently.^5^ In the face of exponential growth in clinically relevant genomic findings driven by precision oncology efforts, there will likely be increased inability for physicians to command the most current information, resulting in increasing delay between academic discovery and clinical integration of variant information. This information gap between the knowledge of variant function and its clinical integration has been described as the “interpretation bottleneck”.^4–6^

Alleviating the interpretation bottleneck will require co-development of targeted sequencing panels and bioinformatic tools that effectively elucidate and annotate clinically actionable variants from sequencing data.^7,8^ These two requirements each raise separate challenges. Commercial and academic pan-cancer clinical gene capture panels have now become commonplace, with at least two obtaining Food and Drug Administration (FDA) approval (FoundationONE CDx^9^ and MSK-IMPACT^10^). Even so, few panels indicate how genomic loci are selected for panel inclusion (**Supplementary Table 1**), and none have proposed a sustainable or scalable mechanism to allow for panel evolution over time in response to knowledge advances in molecular oncology. Variant interpretation perhaps poses even greater and more persistent challenges. Commercial strategies typically rely on the manual curation and organization of research findings into structured databases, which are expensive to create and maintain, forcing companies to limit public access or to charge for use. The resulting lack of transparency creates inefficiencies in the field through unnecessary replication of experiments and suboptimal communication with clinicians, ultimately hindering development of effective patient treatment plans. Separately, governmental and academic institutions have developed variant interpretation resources like COSMIC^11^, ClinVar^12^, and cBioPortal^13,14^ that have drastically improved research efforts and academic discovery, however, these resources do not have well-described clinical relevance summaries for all variants that can be easily accessed and utilized by physicians. Many resources that do provide detailed clinical interpretation of cancer variants (e.g., oncoKB^15^, Cancer Genome Interpreter^16^, Clinical Knowledgebase^17^, and others) but these databases are either limited by license restrictions or closed curation models. The OncoPaD^18^ portal provided the first prospective method to create a rational design for gene panels by linking clinically relevant variants to genomic loci based on a cohort of tumor samples, but it is not linked to detailed and actively updated clinical interpretations.

To address these limitations, here we describe a strategy to integrate a publicly sourced, open variant knowledgebase with an inexpensive and modular platform for targeted gene enrichment and error-corrected sequencing. The Clinical Interpretations of Variants in Cancer (CIViC) database is a freely-accessible, publicly curated repository of therapeutic, prognostic, predisposing, and diagnostic information in precision oncology that permits rapid integration of academic information into an accessible platform for scientists, physicians, and patients^19^ (Supplementary Figure 1). As a proof-of-principle, we utilized this knowledgebase to inform rational design of a targeted sequencing reagent for pan-cancer relevant genetic variants through single-molecule molecular inversion probe (smMIP) capture, which integrates multiplexed targeted sequencing with unique molecular identifier (UMID)-mediated sequence error correction. smMIP capture provides an inexpensive, scalable, and modular platform for targeted gene sequencing, which can be scaled to ultrasensitive levels of variant detection, and which can accommodate the integration of additional probes in response to evolving assay needs. Combined, these two technologies provide a testing framework that links physical genomic loci to relevant clinical information. In this study, we first assess the feasibility of using an open-sourced database to impartially identify clinically relevant variants and design smMIPs probes for their identification. The reagent design was evaluated by comparing CIViC smMIP capture results to orthogonal exome sequencing for 27 cancer samples (Figure 1). Ultimately, we hope that this research will inform development of a clinically validated panel that can dynamically integrate new evidence in precision oncology, enabling systemization of methods for detecting and interpreting cancer variants.

**Figure 1.**
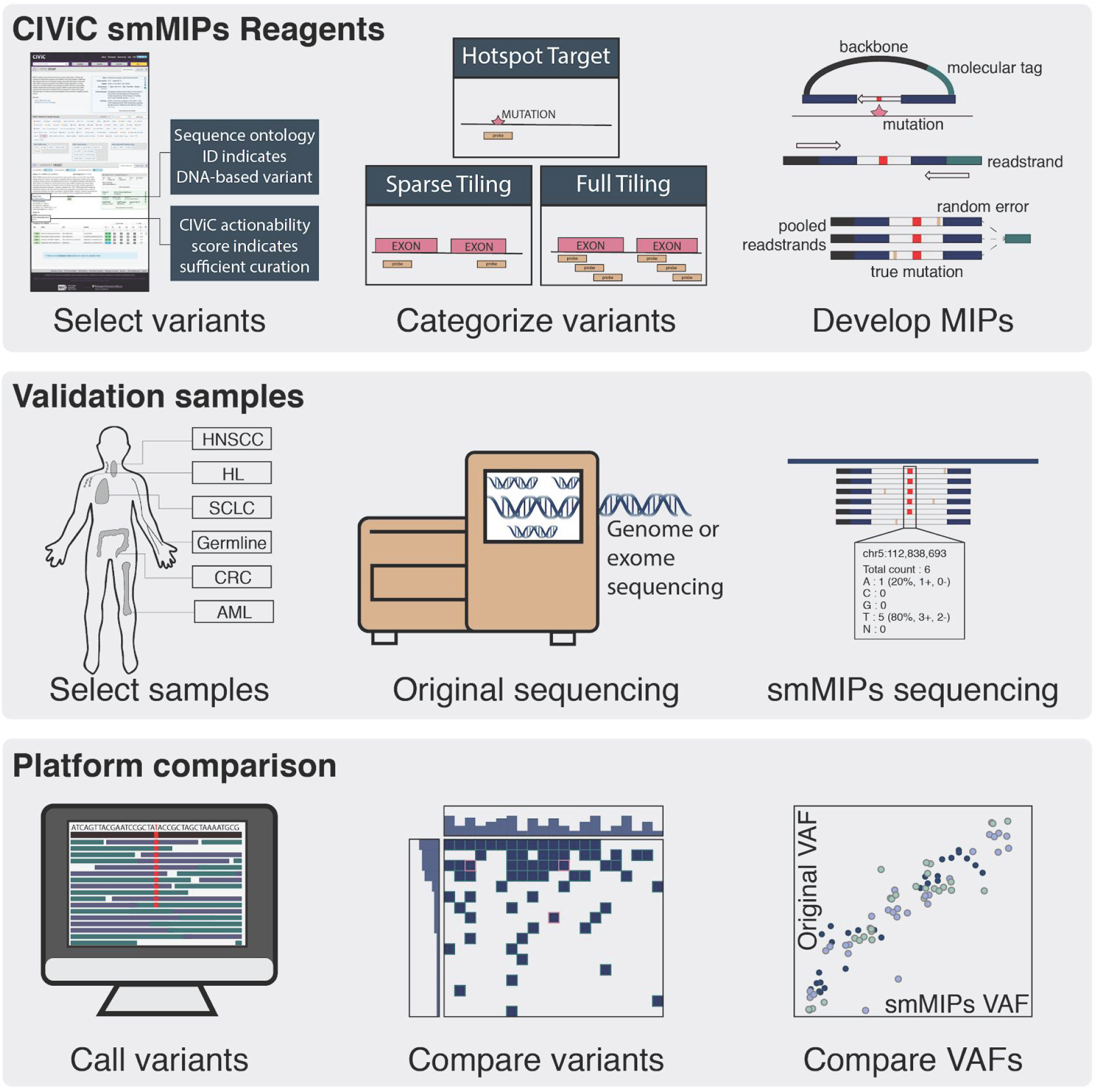
Methods for CIViC smMIPs development and validation. The first series describes CIViC smMIPs development. Variants were selected using the sequence ontology ID and the CIViC actionability score. Subsequently, eligible variants were categorized based on length and smMIPs reagents were designed to target regions of interest. The second series describes sample selection and sequencing methods. In total, there were 22 tumor sample derived from 5 tumor subtypes. Of these 27 samples, 15 had tumor and paired normal samples and 7 were tumor-only samples. The third series shows the analysis used to validate CIViC smMIPs design. Variants were called using the pipeline described in the methods, accuracy was attained by comparing original sequencing variants to CIViC smMIPs sequencing variants, and variant allele frequencies across both platforms were also compared.

## Materials and Methods

### CIViC actionability score development

A CIViC actionability score was created for each variant within the CIViC database to establish whether the existing level of curated evidence warrants inclusion for smMIP targeted capture. The CIViC actionability score determines: 1) the strength of the evidence that was curated and 2) the total level of curation that has been completed for each variant. To determine evidence strength, the evidence level score and the trust rating score were calculated. The evidence level score is a 10-point scale that weighs the evidence strength based on category. Broadly, highest points are awarded to large clinical studies and lower points are awarded to case studies, in vitro studies, and inferential evidence. The trust rating score is a 5-star scale that reflects the curators confidence in the quality of the study within those categories. To determine the total level of curation for each variant, evidence level scores were multiplied by trust rating scores and summed across all evidence items (Supplementary Figure 2). This final value (i.e., the CIViC actionability score) was implemented into the CIViC database using the Variant Application Programming Interface (API) endpoint.

### Determining eligible CIViC variants for smMIP capture

Variants that attained a CIViC actionability score of greater than 20 points were eligible for smMIP targeting. After filtering on the CIViC actionability score, variants were filtered using the curated Sequence Ontology IDs (SOIDs) to only include variants that can be analyzed using a DNA-based sequencing platform. Within CIViC, SOIDs were manually grouped as either: “DNA-based”, “RNA-based”, and/or “Protein-based” (**Supplementary Table 2**). For example, variants with the Variant Type of “missense_variant” would be labeled as “DNA-based,” whereas variants with the Variant Type of “transcript_variant” would be labeled as “RNA-based”. Variants that had a “DNA-based” SOID were eligible for the smMIP targeting and variants whose SOIDs were “RNA-based” and/or “Protein-based” were ineligible. Eligible variants were further filtered if any of the following applied: 1) all evidence supported only germline clinical relevance, 2) evidence was directly conflicting (e.g., one evidence statement detailed sensitivity to a therapeutic and a different evidence statement detailed resistance to the same therapeutic in the same disease setting), or 3) a majority of evidence in a bucket variant (e.g., MUTATION) pointed to a hotspot that was already being covered.

### Designing smMIPs for the CIViC capture reagents

All eligible variants were further categorized by variant length. Using CIViC curated coordinates, variant length was determined (i.e., variant start position minus variant stop position), to determine the number of smMIPs probes required to adequately assess each variant. If the variant length was <250 base pairs, the variant was eligible for hotspot targeting. If the variant was >250 base pairs, the variant required tiling of the protein coding exons. Some large-scale copy number variants (i.e., “AMPLIFICATION”, “LOSS”, “DELETION”), were eligible for sparse tiling, wherein 10 probes distributed across the exons of the gene were retained to enable assessment of copy number state. Other variant types such as “MUTATION”, or “FRAMESHIFT MUTATION”, etc., required tiling of all protein coding exons (**Supplementary Table 3**).

For variants that required hotspot targeting, smMIPs probes were designed for the genomic region indicated in the CIViC database. For all variants that required sparse exon tiling or full exon tiling, the representative transcript from the CIViC database was used to obtain all possible exons associated with each gene from Ensembl (biomart=“ENSEMBL_MART_ENSEMBL”, host=“grch37.ensembl.org”, dataset=“hsapiens_gene_ensembl”). Exons were filtered by Biotype to remove untranslated regions. For variants that required sparse exon tiling, approximately 10 smMIPs were designed to cover a portion of the transcript. For variants that required full exon tiling, overlapping smMIPs (i.e., at least one basepair of overlap) were designed to tile across all protein coding exons in the gene that encompassed the variant. smMIPs were designed and synthesized as previously described with the single alteration that the “-double_tile_strands_separately” flag was used to separately capture each strand of DNA surrounding the target.^20^

### smMIP sequencing and data analytics

Sequencing library construction and balancing of the probe pool were performed as described previously^20^, and sequencing was performed using an Illumina Nextseq 500. Probes were excluded from the final reagent if they demonstrated poor hybridization to target sequence during initial quality checks.

Sequence data analysis was performed as previously described^20^ with three enhancements. First, consensus reads were generated using fgbiotools (http://fulcrumgenomics.github.io/fgbio/) CallMolecularConsensusReads utility with parameters “--error-rate-post-umi=30 --min-reads=2 --min-input-base-quality 20”. Second, a custom variant caller was utilized to identify all consensus calls at a site having at least 2 supporting reads with a minimum specified mapping quality (mapping quality score > 0). Third, variants were required to be detected on at least four DNA strands (at least 2 positive and at least 2 negative) in order to be considered real, rather than post-biological artifact.^21^ Collectively, these provisions require that at least two reads are derived from a common UMID to create a consensus read and that multiple consensus reads in both directions support the apparent variant. This helps to exclude pre-analytic artifacts reflecting DNA damage and stochastic errors that occur during library construction and sequencing.

### Orthogonal sequencing and data analytics

Orthogonal sequencing data from previously conducted whole exome or genome sequencing was used to validate variants identified using CIViC smMIP sequencing. Sequencing alignment and somatic variant calling for the AML31 sample was performed according to *Griffith et al*.^22^ Briefly, reads were aligned to GRCh37 using BWA v0.5.9^23^ and variants were called using one of seven variant callers listed in the manuscript. Sequencing data from the SCLC cases, OSCC cases, and HL cases were analyzed using the Genome Modeling System^2^ at the McDonnell Genome Institute. Reads from these studies were aligned to the reference genome (hg19/GRCh37 or hg38/GRCh38) using BWA-MEM v0.7.10^24^ and duplicates were marked by Picard^25^ and/or SAMBLASTER v0.1.22.^26^. For the SCLC cases, Single nucleotide variants (SNVs) were called using SomaticSniper^27,28^, VarScan^29^, and Strelka^30^; small insertions and deletions (indels) were called using GATK^31^, Pindel^32^, VarScan2^33^, and Strelka. For OSCC cases, SNVs were detected using SomaticSniper v1.0.4, VarScan2 v2.3.6, Strelka v1.0.11, SAMtools r982^34^, and Mutect v1.1.4^35^. Small indels were detected by GATK v5336^36^, VarScan2, Strelka, and Mutect. For HL cases, SNVs were called using the intersection of SomaticSniper v1.0.4, VarScan v2.3.6, Strelka v1.0.11, and Mutect v1.1.4, and indels were called using GATK, Pindel v0.5, VarScan v2.3.6, and Strelka v1.0.11. For these three cohorts, variants identified by automated callers were subjected to heuristic filtering (removal of variants with low VAF [<5%] or low coverage [<20X in tumor or normal track]) and false positives were removed via manual somatic variant refinement.^37^ If coordinates were aligned to GRCh38, final variants were lifted over to GRCh37 using LiftOver.^38^ For the CRC cohort, sequencing, variant calling, and clinical annotation were performed according to methods highlighted in *Pritchard et al.*^39^ Briefly, sequencing was performed using Illumina next-generation sequencing (Illumina, San Diego, CA) and sequencing reads were aligned using BWA v0.6.1-r104 and SAMtools v0.1.18. Indel realignment was then performed using GATK v1.6 and duplicate reads were removed using Picard v1.72. SNV and indel calling was performed through the GATK Universal Genotyper using default parameters and VarScan v2.3.2.

### Rescue and annotation of clinically relevant variants

We compared the variants identified by the CIViC smMIP sequencing pipeline to the variants called by original exome or genome sequencing for samples that had matched tumor and normal sequencing. Variants that were specific to the CIViC smMIP sequencing pipeline were further evaluated for variant support in the original exome or genome sequencing. Using the manual review guidelines outlined in *Barnell et. al.*^*37*^, we manually reviewed genomic loci for smMIPs-only variants using both the smMIPs aligned BAM files and the original exome or genome aligned BAM files. Variants were grouped into four categories based on review of smMIP and original sequencing: 1) germline polymorphism, 2) pipeline artifact, 3) variant support on smMIP sequencing but no variant support on original sequencing, 4) variant support on both smMIP sequencing and original sequencing. smMIPs-only variants were considered germline polymorphisms if there was sufficient support for the variant in the normal track. Variants were considered attributable to pipeline artifacts if aligned reads showed low variant support and/or reads indicated poor mapping/alignment. The remaining variants, regardless of the extent of support in original sequencing, were annotated using curated coordinates in the CIViC database. smMIP-only variants derived from tumor-only samples were further filtered using gnomAD^40^ and annotated using the CIViC database. For variants that showed support on smMIPs sequencing but no variant support on original sequencing, we calculated the likelihood that the variant would be detected using the original coverage and the observed smMIPs VAF. Given that original sequencing required at least 4 reads for a variant to be called somatic, we calculated the binomial probability of obtaining ≤3 variant-supporting reads given the original coverage (number of chances to get a variant supporting read) and the observed smMIPs variant allele frequency (likelihood that a read would show variant support). If the binomial probability of ≤3 variant-supporting reads was >95%, then it was considered statistically unlikely that a variant would be called using original sequencing data.

### Code and accessibility

All raw data, analysis, and preprocessing code, are publically available on the GitHub repository. All plots were produced using the MatPlotlib library in Python.^41^ The raw sequencing data are publically available for most projects included in this study (**Supplementary Table 4**). The smMIP sequence analysis pipeline is accessible on bitbucket.

### URLs

Clinical Interpretation of Variants in Cancer database, http://www.civicdb.org/. CIViC Interface public API, http://griffithlab.org/civic-api-docs/. CIViC Panels API https://civicdb.org/api/panels. GitHub, https://github.com/. smMIPs GitHub Repo, https://github.com/griffithlab/civic-panel/tree/master/smMIPs_Manuscript. Ensembl GRCH38 BioMart, http://useast.ensembl.org/biomart/martview/c2a89c09bd76058dfd5a9821453a1da7. smMIP sequence analysis pipeline, https://bitbucket.org/uwlabmed/smmips_analysis.

## Results

### Identification of eligible CIViC variants for smMIP targeting

At the time of CIViC smMIPs reagent design, there were 988 variants spanning 275 genes within the CIViC database that had at least one evidence item. After filtering based on CIViC actionability score and SOID (see **Methods**), smMIPs were designed to cover all eligible CIViC variants. A set of 2,097 probes were developed and tested on control samples. Of these, 70 probes showed poor capture efficiency and were subsequently eliminated from the analysis. Removal of the underperforming probes affected 32 variants across 16 genes. The final capture reagent targeted 111 CIViC variants spanning approximately 61.5 kb of genomic space (**Supplementary Table 3**). Of these variants targeted by smMIP capture, 71 required hotspot targeting, 14 variants required sparse exon tiling, and 26 required full exon tiling. The 111 variants covered by CIViC smMIP sequencing were based on 1,168 clinically relevant evidence items whereby 820 (70%) evidence items predicted response to a therapeutic, 232 (20%) detailed prognostic information, 52 (4%) indicated diagnostic information, and 64 (6%) evidence items supported predisposition to cancer (Figure 2).

**Figure 2.**
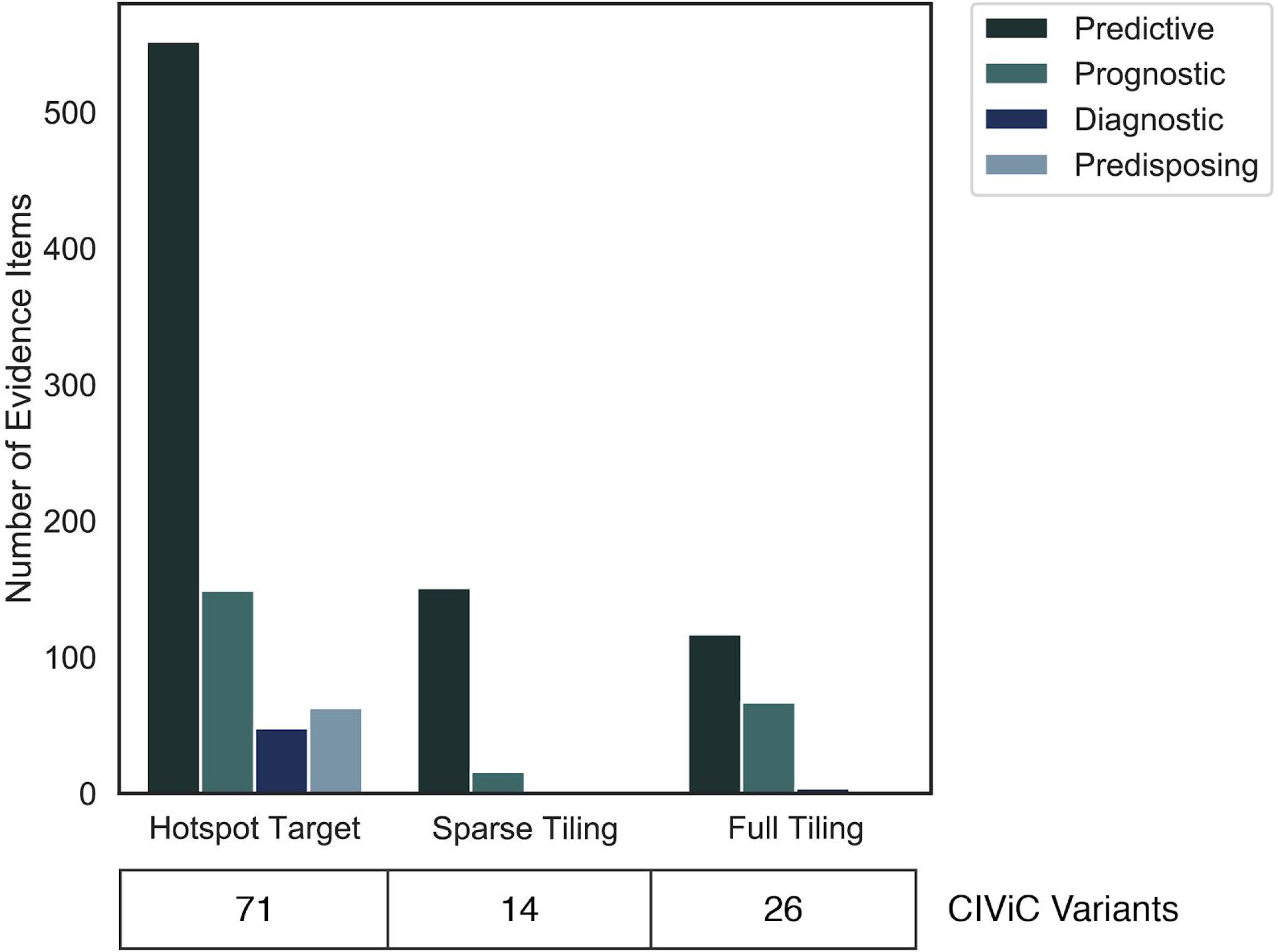
Regions targeted by the CIViC single molecule molecular inversion probes (smMIPs) are, by design, supported by extensive clinical relevance according to the CIViC database. Variants that were eligible for CIViC smMIPs development were bucketed into various coverage methods based on sequence ontology identification number (SOID) and length. Bar graph shows total number of evidence items used for each of the buckets parsed by the evidence type.

After reagent development, the CIViC database was re-evaluated to determine additional variants that did not meet the original actionability score threshold, but were encapsulated by the designed smMIPs. The total number of variants covered by the CIViC smMIPs, regardless of actionability score, was 408 variants spanning 49 genes. The majority of these additional variants (n = 203) were contained within VHL, an area of very active CIViC curation.

### Tumor samples used to validate CIViC smMIPs design

We assembled validation samples from 5 different cancer genomic studies. Four studies were conducted at the McDonnell Genome Institute at Washington University School of Medicine and one study was performed at the University of Washington (**Supplementary Table 4**). In total, we obtained tumor and paired normal samples from 5 individuals with head and neck squamous cell carcinoma (HNSCC), 9 individuals with small cell lung cancer (SCLC)^42^, and 1 individual with Hodgkin’s lymphoma (HL). We also obtained tumor-only samples from 1 individual with HL, 1 individual with acute myeloid leukemia (AML)^22^, and 5 individuals with colorectal cancer (CRC). In total, there were 37 samples evaluated from 22 individuals.

Each of the 22 individuals had previously undergone exome or genome sequencing, automated somatic variant calling, and somatic variant refinement via manual review (see **Methods**). Using exome or genome sequencing the total number of putative somatic variants called for these 22 samples was 12,602. The average variant burden was 573 variants per sample with a range of 2 to 3,900 variants per sample. The variant coordinates from these samples were compared to the genomic region covered by the CIViC smMIP capture to determine potential validating variants. In total, there were 84 variants identified via exome or genome sequencing that overlapped with the CIViC smMIPs region assayed (**Supplementary Table 5**).

### smMIP sequencing and data analysis

#### Initial quality check

The average number of total tags captured for all samples was 5.4 million (standard deviation = 3.3 million tags). Two HNSCC tumor samples had significantly fewer reads than the rest (i.e., greater than 1 standard deviation from the mean) and one HL sample had reduced tag complexity relative to the rest (i.e., fewer than 600,000 unique captured mips). These results are consistent with poor template quality and/or quantity, and the samples were excluded from the analysis. There were seven normal samples that had either fewer reads or low complexity due to low input mass (100-200 ng). Post quality check, 33 samples derived from 19 individuals were eligible for reagent validation. These samples had 65 variants derived from orthogonal sequencing that had overlap with the CIViC smMIPs coverage (Figure 3). The average consensus read depth for these 65 variants was 2,942 reads (std = 4,697 reads).

**Figure 3.**
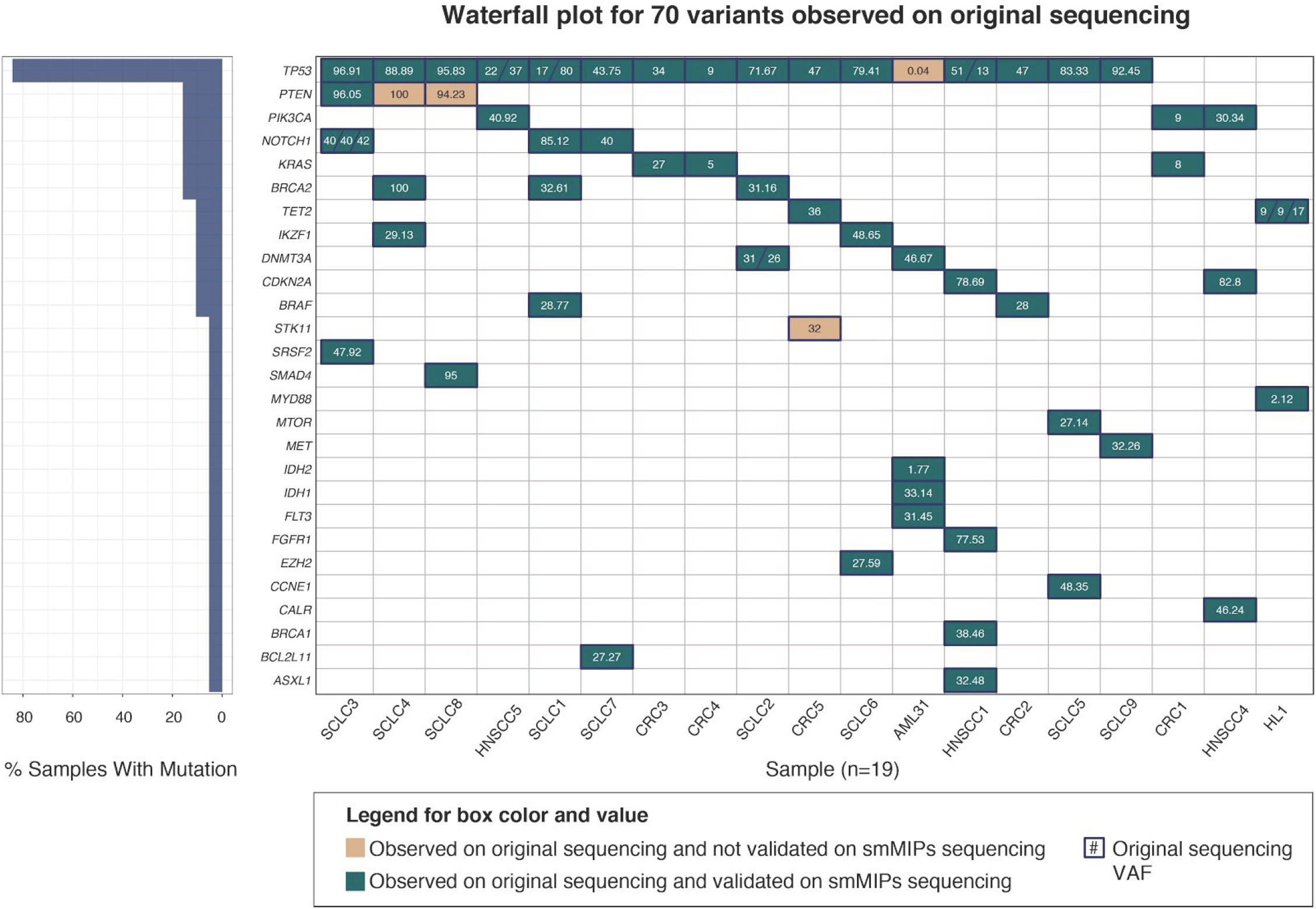
Waterfall plot shows extensive overlap between variants observed using original exome or genome sequencing with variants observed using CIViC smMIPs sequencing. Each column represents a sample that had original exome or genome sequencing with subsequent orthogonal validation using the CIViC smMIPs sequencing. Rows represent mutated genes across all samples. Numbers within each box represent the variant allele frequency (VAF) observed on original exome or genome sequencing. Green boxes indicate that a variant was observed by CIViC smMIPs and validated with original exome or genome sequencing. Tan boxes indicate the the variant was observed on original exome or genome sequencing but not identified via the CIViC smMIPs sequencing. The left panel indicates the number of samples containing a mutation in the indicated gene.

#### Accuracy of CIViC smMIP variant identification compared to exome or genome variant identification

Of the 65 variants identified on exome sequencing, all but 4 were also identified using CIViC smMIP sequencing (Figure 3). One variant was missed due to lack of adequate coverage, two variants were missed due to low performing probes, and one variant was retrospectively considered ineligible due to smMIPs design. The variant missed due to inadequate coverage was a TP53 (p.G266R) variant on AML31. Original sequencing indicated that this variant was present at 0.04% VAF, therefore, given smMIPs coverage of 2,388 reads at this site, there was only a 0.01% chance that this variant would have been detected (one-tailed probability of exactly, or greater than, 4 reads (K) out of 2,388 reads (n); p = .004603). However, this low-prevalence variant could have been recovered given additional sequence coverage. Additionally, there were two variants missed due to low MIP performance. The first variant that was missed (PTEN - C71Y on SCLC8 at 94% VAF) was due to poor performance of the MIP covering the region of interest in the reverse direction. This MIP showed only 1 aligned read across all 36 samples and had no aligned reads in SCLC8. Despite the fact that there was extensive support from the forward MIP (95% VAF with 34 / 35 consensus reads), the requirement that both forward and reverse reads show support prevented this variant from being called (see **Methods**). The second missed variant (PTEN - e8-1 on SCLC4 at 100% VAF) was due to low performance of MIPs in both directions. Even though both the forward and the reverse MIP showed variant support, the forward MIP only contained 2 consensus reads and the reverse MIP only contained 1 consensus read, preventing it from being called as somatic. The final variant (STK - T328N on CRC5 at 32% VAF) was retrospectively considered ineligible because the original smMIPs developed to cover the eligible STK variant called for sparse tiling (i.e., identification of copy number change). As such, the variant was contained by a region that did not have full coverage in the forward direction. When evaluating the reverse MIP that contained this site, we observed a 34% VAF (402 / 1,184 reads), which was comparable to original sequencing. However, lack of a secondary probe designed against the complementary DNA strand prevented this variant from being called as somatic. After removing this variant from the list of eligible variants, the CIVIC smMIP capture sequencing attained a 95% sensitivity for variant detection (n = 64 variants).

#### VAF correlation between CIViC smMIPs sequencing and exome or genome sequencing

Variant allele frequencies (VAF) obtained via exome or genome sequencing were compared to the VAF obtained using the CIViC smMIPs. To compare VAF quantitation across platforms, the 19 variants obtained from samples that failed CIViC smMIPs sequencing quality check were eliminated (Figure 4a). Subsequently, we eliminated the four variants that were not validated using the CIViC smMIPs reagents (Figure 4b). When comparing original VAF to CIViC smMIPs VAFs, Pearson correlation for the remaining 61 variants was 0.885. There were several variants whereby the VAF observed on the CIViC smMIPs sequencing was lower than that observed on the original sequencing. These outliers were not associated with tumor type, sequencing mass input, average coverage, presence of matched normal, or sample type (Figure 4c-g).

**Figure 4.**
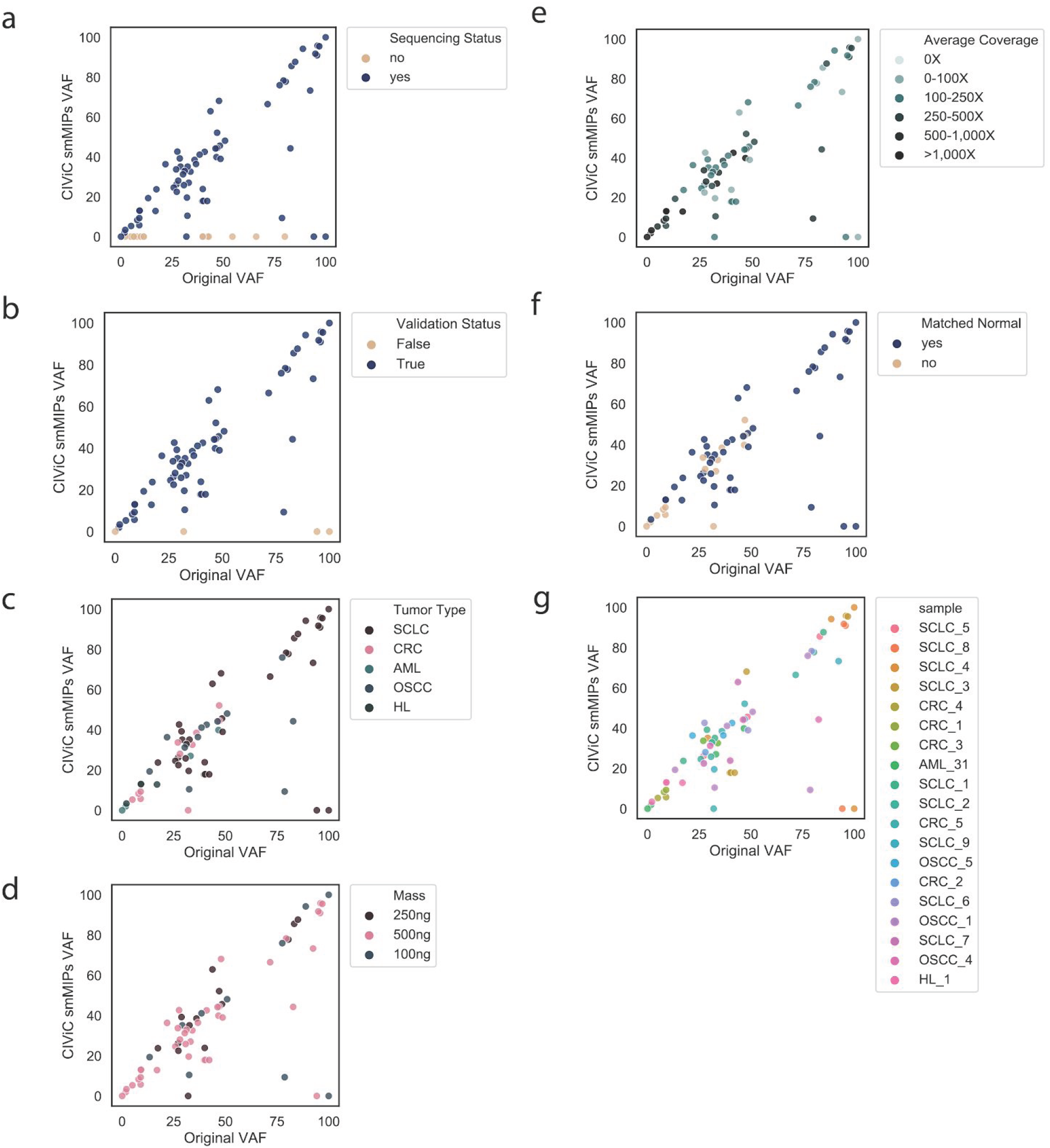
Variant allele frequencies (VAFs) observed using original exome or genome sequencing correlates with VAFs observed using CIViC smMIPs sequencing. **a** Correlation of VAF with original sequencing parse by sequencing status (i.e., passed sequencing if total sequencing counts > 1 standard deviation from mean and tag complexity > 600,000). **b** Correlation of VAF with validation status (e.g., ‘True’ if the variant identified using exome/genome sequencing was identified on CIViC smMIPs sequencing). **c** Correlation of VAF parsed by tumor type. **d** Correlation of VAF parsed by DNA mass input for library construction. **e** Correlation of VAF parsed by depth of sequence coverage. **f** Correlation of VAF parsed by presence or absence of matched normal tissue. **g** Correlation of VAF parsed by sample identity.

### Analysis of variants only identified using CIViC smMIP sequencing

#### CIViC smMIPs capture rescues clinically relevant variants

Using samples that had sequencing data for both tumor and matched normal (n = 12 samples), we evaluated whether the targeted CIViC smMIP sequencing could identify clinically relevant variants that had been missed during the original sequencing annotation pipeline. There were 273 variants recovered by CIViC smMIP sequencing that were not identified using exome or genome sequencing. After manually reviewing these variants within the original exome or genome alignments, 55 variants (20.1%) were identified as germline mutations. smMIP sequencing VAF distribution at 50% and 100% further supported that these variants were germline polymorphisms (Figure 5a). An additional 36 variants (13.2%) were thought to be caused by pipeline artifacts and attributable to assumptions underlying automated callers or alignment problems. The majority of these artifacts were caused by nucleotide repeats in the reference sequence (Figure 5b). There were 171 (62.6%) variants called as somatic using CIViC smMIPs that did not have any variant support on the original sequencing. For these variants, we calculated the binomial probability that ≤3 reads would support the variant given the original coverage (number of chances to get a variant supporting read) and the observed smMIPs variant allele frequency (likelihood that a read would show variant support). If the binomial probability of ≤3 variant-supporting reads was >95%, then it was considered statistically unlikely that a variant would be called using original sequencing data. Using this calculation, 162 variants (94.7%) showed insufficient coverage in the original exome or genome sequencing for detection (Figure 5c). Finally, 11 variants (4.2%) were not called as somatic on original sequencing but did show some variant support in those original sequencing data. The VAFs observed on original sequencing data were strongly correlated with the VAFs observed using CIViC smMIP sequencingl (Pearson r = 0.92) (Figure 5d). Reviewing manual review files from the original sequencing, we observed that 6 of these variants failed manual review due to low VAF, 4 variants had not been called by automated somatic variant callers, and 1 variant failed manual review due to a perceived sequencing artifact.

**Figure 5.**
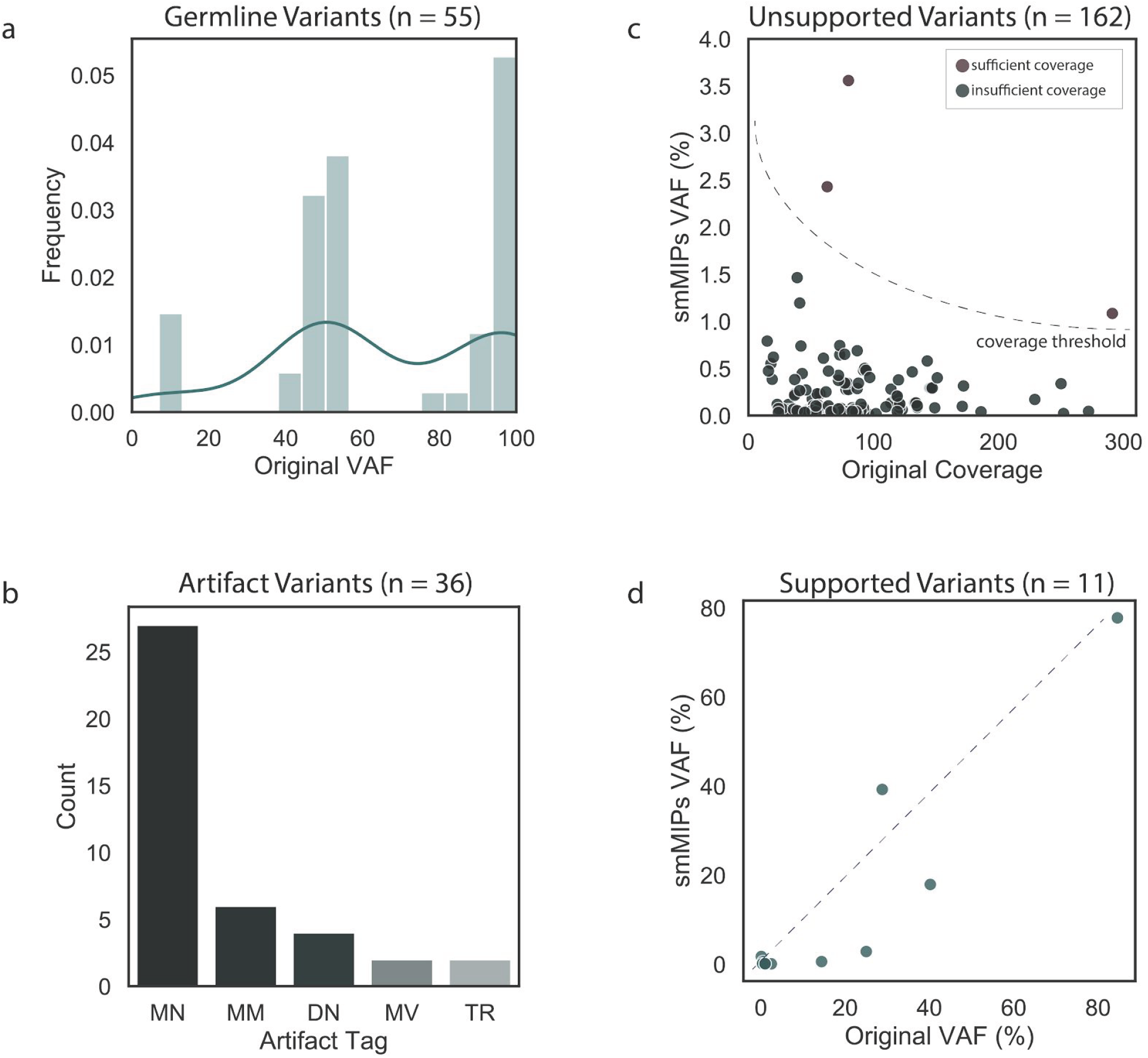
Analysis of variants rescued by CIViC smMIPs sequencing for samples with both tumor and matched normal. There were 217 variants called as somatic by the CIViC smMIPs sequencing that were not identified on original sequencing. All variants were manually reviewed using both CIViC smMIPs sequencing data and original sequencing data. **a.** During manual review 55 variants were identified as germline. A histogram shows that the distribution of the smMIPs VAF for these germline variants are observed at 50% and 100% VAF, indicating heterozygosity and homozygosity, respectively. **b.** An additional 36 variants were identified as sequencing artifacts. Most artifacts were either mononucleotide repeats (MN), dinucleotide repeats (DN), or tandem repeats (TR). Other artifacts include multiple mismatches (MM) or multiple variants (MV). **c.** During manual review, 162 variants did not show any support in the original sequencing data. Most unsupported variants did not have sufficient coverage to be detected based on a binomial probability of at least 4 variant-supporting reads (see **Methods**). **d.** The remaining 11 variants had variant support in original sequencing but were not called as somatic in final original annotation. Scatter plot shows correlation between original VAF and CIViC smMIPs VAF for these variants.

We elucidated the clinical relevance of the 182 variants that were missed by original sequencing but legitimately called as somatic by CIViC smMIP sequencing (i.e., not germline or caused by pipeline artifact). In total, there were 73 CIViC evidence statements associated with variants in the 12 samples that had both tumor and matched normal sequencing data (**Supplementary Table 6**). Of these, 33 were predictive, 4 were diagnostic, 36 were prognostic, and 0 were predisposing.

#### Annotation of CIViC smMIPs variants using tumor-only samples

We also examined the clinical relevance of the 401 variants in tumor-only samples that were called as somatic by CIViC smMIP sequencing. In total, there were 130 CIViC evidence statements associated with the 6 samples that had tumor-only sequencing data. For these 130 CIViC evidence statements, 63 were predictive, 1 was diagnostic, 66 were prognostic, and 0 were predisposing.

## Discussion

The method employed to build a preliminary CIViC smMIPs targeted sequencing reagent offers several advantages relative to existing design paradigms. Use of an open-sourced database provides an unbiased mechanism to survey existing literature within precision oncology to identify variants that are relevant for capture. Additionally, the public CIViC API permits rapid mapping of identified somatic and germline variants to CIViC clinical relevance summaries. Most importantly, the variants covered by CIViC and associated clinical summaries can be updated in real-time as knowledge is entered into the database to accommodate new information discovered within the field of precision oncology. The choice to use the smMIP capture method for sequencing provides inherent error correction capability, scalability to detect ultrasensitive variation, and cost effectiveness within a modular design. Combining the public access CIViC database with an ultrasensitive and versatile capture reagent thus provides an advantageous and principled method for building precision oncology capture reagents. This approach could enable a standardized framework for detecting and interpreting cancer-relevant genomic variation, lowering barriers to use of genomic analysis in the clinical practice of oncology.

The CIViC actionability score was used to obtain a community consensus on the clinical relevance for all variants within the database and the publically curated coordinates were used to design the probes required for variant capture. By comparing variants identified using exome or genome sequencing with variants identified using CIViC smMIPs sequencing, we concluded that CIViC variants and variant coordinates were being accurately incorporated into the database. Overall, the design attained an accuracy of 95% (n = 64 variants) with Pearson r correlation of 0.885 for VAFs of variants identified by both sequencing methods. This finding validates the technological approach of the smMIPs design and that coordinates derived from the CIViC knowledgebase can be used to accurately cover desired variants of interest.

This study used the CIViC actionability score to determine if a variant had sufficient evidence to influence clinical decisions. Filtering out variants with an actionability score < 20 points ensured that smMIPs were designed to only target loci that had clinical implications if mutated. In addition to defining relative level of variant curation, the CIViC actionability score can also be used identify gaps within the database. For example, if an externally validated variant (e.g., IDH1 - R132S) has a low score, users can search the queue of suggested publications in CIViC or find an article to add an evidence statement. Additional curation would subsequently increase the actionability score to better reflect the clinical value of the variant. Addition of the CIViC actionability score to the interface should promote efficient use of curation resources, filling knowledge gaps, and helping to understand when variants have achieved adequate curation coverage.

There were only three orthogonally validated variants that were not identified using CIViC smMIP sequencing. A subclonal TP53 mutation from AML31 was not identified due to low tumor VAF Originally, this variant was not observed on exome sequencing at 160X, genome sequencing at 314X, or capture sequencing at 1,167X; however, 3 variant supporting reads (0.04%VAF) were detected using a capture panel with spike-in probes for recurrently mutated AML variants (6,117X).^22^ Even though this variant was not identified by the CIViC smMIP sequencing reagents at a sequencing depth of 2,388, it is likely that increased sequencing depth at this locus would have resulted in variant detection. This is further supported by the fact that other variants with low VAFs (e.g., VAF = 1.77%) were readily identified by the CIViC smMIP reagent. The PTEN variants on SCLC8 and SCLC4 were missed because one or more probes that covered the region of interest showed reproducibly poor performance across samples. Improved balancing of smMIP covering this region, redesign of failed probes, or denser tiling of poor performing regions could improve performance in a subsequent design.

The STK p.T328N variant on CRC5 is noteworthy because it reflects the dynamic nature of academic discovery and illustrates the potential advantage of a panel that permits evolution in response to changing literature. This variant was incidentally found because the original CIViC smMIP capture design was created when STK-LOSS was the only clinically relevant STK variant that passed the CIViC actionability score threshold. Given the “LOSS” variant type, the variant only required sparse tiling and approximately 10 MIPs were designed to capture the variant. In July of 2018, Skoulidis et. al.^43^ published an article demonstrating that STK mutations can cause anti-PD-1 resistance. This article was entered into the CIViC database in August of 2018 under STK - MUTATION. Curation on this variant raised the CIViC actionability score for STK - MUTATION to >20 points. Therefore, if the reagents had been designed after these findings, STK - MUTATION would have been eligible for CIViC smMIPs design. As such, STK-MUTATION would have been bucketed into the full-tiling cohort, MIPs covering this particular STK variant would have been designed, and the variant would have a CIViC annotation of, “resistance to Pembrolizumab, Nivolumab, and Atezolizumab”. This anecdote illustrates the advantages of a dynamic capture panel that can incorporate new discoveries in real-time.

Like all targeted reagents, the preliminary CIViC smMIP design has certain limitations that can be addressed with future iterations. First, the reagent design is limited by the current curation within the CIViC database. Extensive curation from certain groups (e.g., the University Health Network has focused on curating VHL variants) disproportionately increases representation. Conversely, lack of curation in certain areas show a disproportionate decreased representation. Intersection of the CIViC actionability score with existing databases can identify gaps within the database to highlight the need for external and internal curation. Second, targeting additional variants that become eligible as the literature evolves requires designing new probes and rebalancing the reagent. Integrating software to design probes and estimate required probe proportions into CIViC could help automate this process for future iterations. We also observed in this study that some samples failed quality check due to low sample quality, which potentially limits this approach for degraded samples or samples subjected to formalin-fixed paraffin-embedded (FFPE) preservation, which could potentially be improved with changes to specimen processing or capture procedures. Finally, intersection of variants covered by CIViC smMIPs reagents with clinical relevance summaries from CIViC requires use of the CIViC API, which might be a barrier for some individuals. Future studies will improve visualization of clinical relevance summaries associated with identified variants. Ultimately, after addressing these limitations, we hope to develop, optimize, and validate a clinical CIViC smMIP oncology panel according to the guidelines provided by *Jennings et al*.^44^ Using the methods provided herein, we plan to build and test panel reagents for a clinical-grade assay validated by current best practices.

In summary, this research validates that community curated data on clinically relevant cancer variants provides an unbiased and dynamic method for capture reagent design. The curated coordinates in the database accurately map to desired variants and probes designed using these coordinates show accurate recapitulation of the genomic landscape described by orthogonal sequencing.

## Acknowledgements

We gratefully acknowledge Tim Ley for sharing genomic data that made this project possible. We also would like to thank the patients and their families for their selfless contribution to the advancement of science. EKB was supported by the National Cancer Institute (T32GM007200 and U01CA209936). Select sample data was funded by the Genomics of AML PPG (T. Ley, PI, P01 CA101937). RU was funded by the National Comprehensive Cancer Network Oncology Research Program, general research support provided by Novartis Pharmaceutical Corporation (Novartis), the V Foundation for Cancer Research, and the National Institute of Dental and Craniofacial Research (NIH NIDCR R01DE024403). RG is funded by the National Cancer Institute (NIH NCI U01CA231844). MG is funded by the National Human Genome Research Institute (NIH NHGRI R00HG007940). SJS is funded by the National Cancer Institute (NIH NCI R33CA222344). OLG is funded by the V Foundation for Cancer Research, and the National Cancer Institute (NIH NCI U01CA209936 and NIH NCI U01CA231844). This research was supported by a Cancer Moonshot funding opportunity, specifically, an Activities to Promote Technology Research Collaborations (APTRC) for Cancer Research (Admin Supp) award to SJS and OLG under parent awards (R33CA222344 and U01CA209936).

## Author Contributions

EKB completed reagent development, clinical data analysis, wrote code, and wrote the manuscript. KP conducted sequencing analysis. SJS and AW developed and validated the smMIPs reagents, provided WES data and tumor tissue, and wrote the manuscript. KMC created figures and wrote the manuscript. TAF, RU, and RG, provided WES data and tumor tissue and edited the manuscript. MG, SJS, and OLG supervised the project and revised the paper.

## Supplementary Tables and Figures

### Supplementary Tables

https://docs.google.com/spreadsheets/d/1iYc-EYtNNMd9eSqeKZqKU0TQMdmP6aZJ4ZS3oF3BW5s/edit?usp=sharing

**Supplementary Table 1. List of existing pan-cancer capture reagents with description on genes and development.**

**Supplementary Table 2. Sequence ontology identification numbers (SOIDs) with sequencing category.**

**Supplementary Table 3. Variants and associated genes eligible for CIViC smMIPs design after filtering based on CIViC actionability score and SOID.**

**Supplementary Table 4. Sequencing data availability for samples used in analysis.**

**Supplementary Table 5. Samples used to validate the CIViC smMIPs reagents with variants and variant allele frequencies identified using whole exome sequencing.**

**Supplementary Table 6. The CIViC database was used to elucidate clinical relevance for the 218 variants missed by original sequencing. For each of the 12 samples that had matched tumor/normal pairs, the clinical relevance statement, associated evidence item ID, and the pubmed ID are listed.**

### Supplementary Figures

**Supplementary Figure 1.**
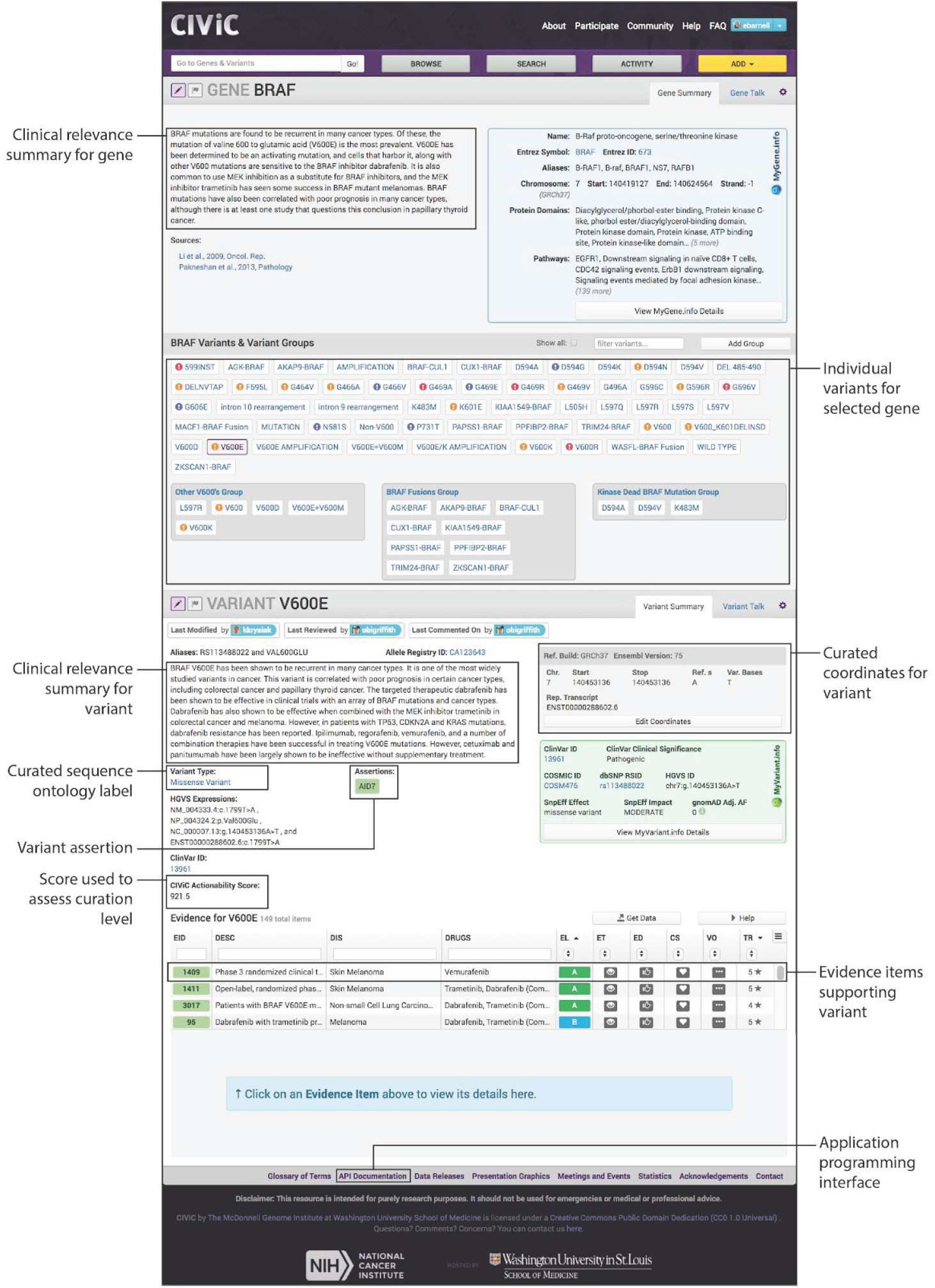
Example of CIViC interface for BRAF V600E mutation with highlighted features. The clinical relevance summary provides an overview of the clinical relevance of the gene. Individual variants and variant groups associated with the gene are subsequently listed for selection. Selecting a variant (i.e., V600E) allows the user to see the clinical relevance summary for the variant and additional curated information (i.e., sequence ontology label, variant coordinates, clinical assertion, and CIViC actionability score, etc.). Evidence statements supporting the variant can be viewed by clicking on the rows in the evidence statement grid. This provides the user with additional information on how the clinical relevance statements and actionability score were generated.

**Supplementary Figure 2.**
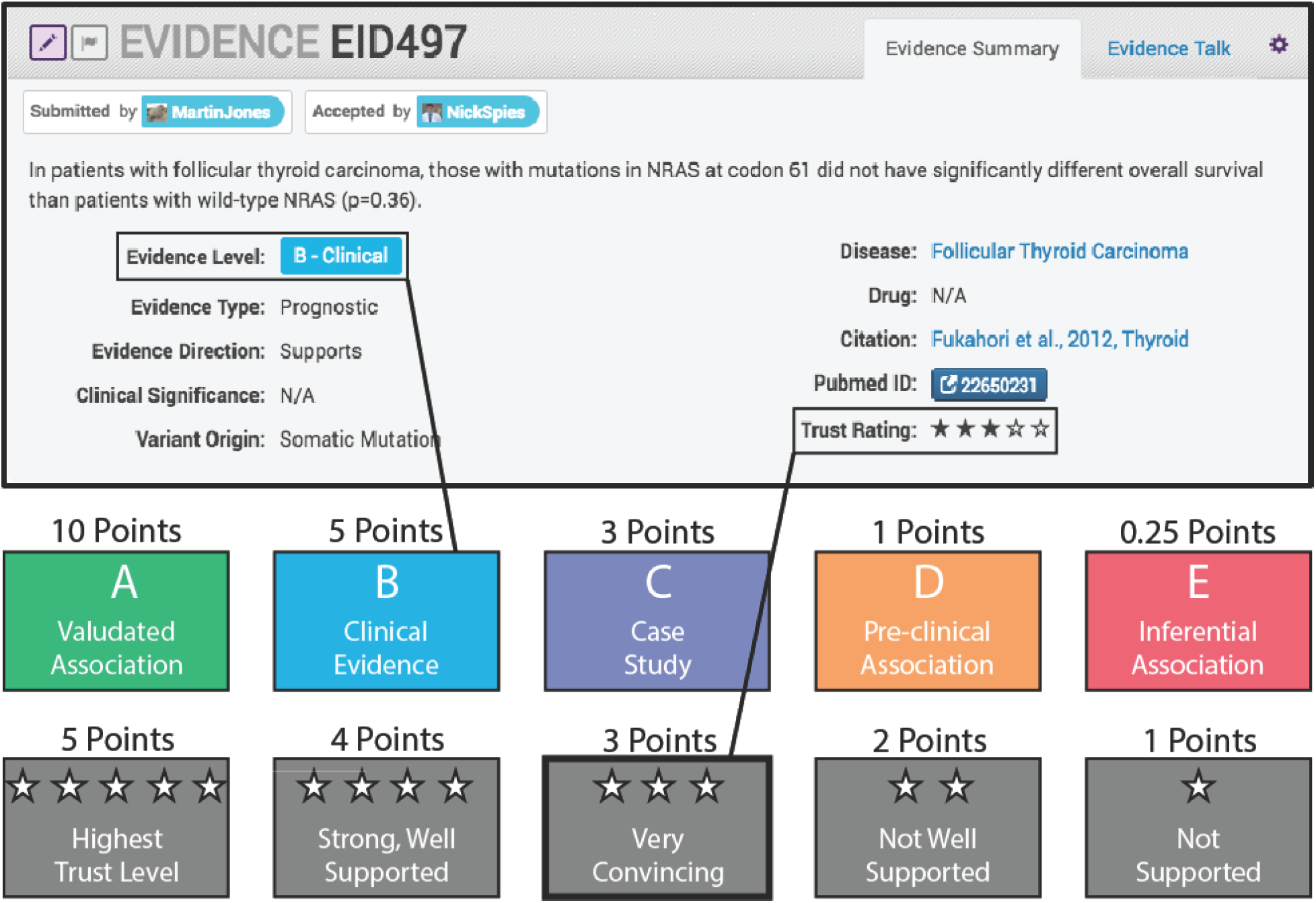
The CIViC actionability score provides relative level of curation for variants within the CIViC database. To calculate the actionability score, first evidence item scores are calculated by multiplying the associated evidence level point by the trust rating point. For each variant, evidence item scores are summed and the CIViC actionability score is generated.

